# Age-related auditory nerve deficits propagate central gain throughout the auditory system: Associations with cortical microstructure and speech recognition

**DOI:** 10.1101/2025.05.19.654964

**Authors:** Emily M. Fabrizio-Stover, James W. Dias, Carolyn M. McClaskey, Kelly C. Harris

## Abstract

There is growing evidence that many perceptual difficulties associated with age-related hearing loss are not solely due to cochlear damage and are exacerbated by changes within the central nervous system. We examined electrophysiological (EEG) responses to clicks and diffusion kurtosis imaging (DKI) in 49 older (29 female) and 26 younger (20 female) adults to determine the extent to which auditory nerve (AN) deficits in older adults contributed to functional and structural changes throughout the auditory system. Older adults exhibited smaller AN responses, similar brainstem responses, and larger auditory cortex (AC) responses, demonstrating progressive “central gain”. Audiometric thresholds were not predictive of EEG measures. Reduced AN function predicted deficits in cortical microstructure (lower AC fractional anisotropy, FA) in older adults, consistent with myelin degeneration. These lower FA values in the AC of older adults also predicted larger AC responses and more central gain. Older adults exhibited significantly lower AC FA and higher mean diffusivity (MD) than younger adults, and AC FA and MD were significant predictors of speech-in-noise (SIN) recognition in older adults. The results suggest that reduced afferent input in older adults not only results in functional changes throughout the auditory system consistent with progressive gain, but also contributes to deficits in AC structure beyond those explained by age alone, contributing to SIN deficits. Understanding the complex effects of age, reduced AN input, central gain, and AC structure on SIN recognition may provide potential therapeutic targets for intervention.

**Significance:** Age-related hearing loss is the most common sensory deficit with aging, but little is known about how the auditory cortex adapts to the chronic loss of afferent auditory input. We measured auditory nerve, brainstem, and cortical responses in younger and older adults. In older adults, deficits in afferent input predicted progressive hyperexcitability, or “central gain”, at the brainstem and cortex. Smaller auditory nerve responses predicted poorer white matter structure in auditory cortex. Poorer white matter structure predicted smaller auditory cortex responses, more auditory cortical gain, and worse speech-in-noise recognition in older adults. This is the first study to show that reduced afferent input leads to both amplified cortical responses and reduced cortical integrity in older adults, contributing to speech-in-noise deficits.

## Introduction

Older adults with and without hearing loss often experience difficulties recognizing speech, particularly in challenging listening environments (Dubno et al., 1984; Başkent et al., 2014; Goossens et al., 2017; Holder et al., 2018; Kocabay et al., 2022). Many studies of aging and speech recognition focus on characterizing the auditory periphery (Zeng et al., 2005; Hoben et al., 2017; Bramhall and McMillan, 2024). Although age-related hearing loss, and even more so auditory nerve (AN) deficits (Makary et al., 2011; Dias et al., 2024), are one of the most common conditions of aging, little is known regarding how the central auditory system adapts to this progressive decline in auditory input.

Our prior research (Harris et al., 2022; Rumschlag et al., 2022) and others (Salvi et al., 2017; Johannesen and Lopez-Poveda, 2021; Hutchison et al., 2023) suggest that in older adults, the loss of afferent input contributes to hyperexcitability, or “central gain”, in the auditory system that may be maladaptive to speech-in-noise (SIN) recognition.

These studies demonstrate that central gain changes are detectable early in the auditory pathway and are maintained or potentially further increased at higher (i.e., cortical) levels of processing. Heterogeneous effects are reported following transient auditory deprivation in younger adults, with central gain occurring in the brainstem (Maslin et al., 2013; Brotherton et al., 2017), the auditory cortex (AC) (Graterón et al., 2022), or both (Hutchison et al., 2023). Moreover, plasticity associated with chronic conditions, like age-related deafferentation, may not be well modeled by such transient manipulations (Seybold et al., 2012). Here, we expanded upon our prior work to examine how age-related AN deficits predict the progression of central gain in the brainstem and AC. We predicted that older adults would exhibit systemic progression of central gain beginning at the brainstem with greater central gain observed in the AC.

Additionally, myelin integrity is essential for the efficient and accurate encoding of complex stimuli, like speech, especially in difficult listening environments (Zeng et al., 2005). To examine the extent to which afferent loss predicted structural deficits in the AC, we computed metrics of microstructural integrity from diffusion kurtosis imaging (DKI) that are closely associated with myelin. Myelin degeneration plays a fundamental role in neurodegenerative disease and aging (Rawji et al., 2023) and may contribute to cortical hyperexcitability (Borges et al., 2023). Myelin is crucial for the encoding of temporal cues, important for speech recognition. Age-related deficits in auditory temporal processing, consistent with myelin degeneration, are a hallmark of aging (Strouse et al., 1998; Gallun et al., 2014; Ozmeral et al., 2016). However, the role of afferent loss in myelin degeneration and central gain, and the role of these factors in SIN recognition, remain largely unknown in older adults. Previously, a meta-analysis showed that hearing loss is associated with changes in cortical diffusivity throughout the central auditory system, but few studies were analyzed and most had modest sample sizes (Tarabichi et al., 2018). Despite the prevalence of AN deficits in aging, often preceding elevated thresholds (Sergeyenko et al., 2013; Kobrina et al., 2020), no studies have examined the degree to which age-related AN functional deficits predict cortical microstructure and the potential impact of such microstructural deficits on auditory function and central gain.

This is the first study to use a within-subject design to characterize the progression of central gain from the AN to the brainstem and AC in older adults. We also examined how age and decreased afferent input predict diffusion metrics associated with myelin integrity in the central auditory system. We tested the overarching hypothesis that age-related loss of afferent input predicts myelin degeneration in the AC, contributing in part to central gain and poorer SIN recognition. Understanding how a loss of afferent input with age may affect AC microstructure and, in turn, how AC microstructure may affect SIN, can shed light on new therapeutic targets to improve the speech recognition of older adults.

## Materials and Methods

### Participants

Forty-nine older adults [Age: 55-82 years, mean age: 66, SD= 7.870, 29 female] and 26 younger adults [Age: 18-30 years, mean age: 24, SD= 2.942, 20 female] were recruited from the Charleston, South Carolina, community. All participants were native English speakers and had a Mini-Mental Status Examination score of at least 27. Participants with a history of head trauma, seizures, conductive hearing loss or otologic disease, self-reported central nervous system disorders, or an inability to safely undergo magnetic resonance imaging (MRI) were excluded. Younger participants had pure tone audiometric thresholds ≤25 dB HL from 0.25 kHz to 8.0 kHz. Older participants were included if their thresholds were ≤65 dB HL from .25 to 4.0 kHz. No participants had asymmetries between ears in excess of 15 dB HL. Participant audiograms are shown in Figure 1. A pure-tone average (PTA) was calculated for each participant, representing the average hearing thresholds in the test (right) ear from 0.5 kHz to 4.0 kHz. PTAs in older adults ranged from 3.75 dB HL to 46.88 dB HL [Mean= 19.9, SD= 9.987]. PTA was used to examine the potential effects of hearing level on AN function, cortical microstructure, and SIN. PTA was significantly higher in older adults than younger adults [p<0.001]. Participants provided written informed consent before participating in this study approved by the Medical University of South Carolina.

**Figure 1:**
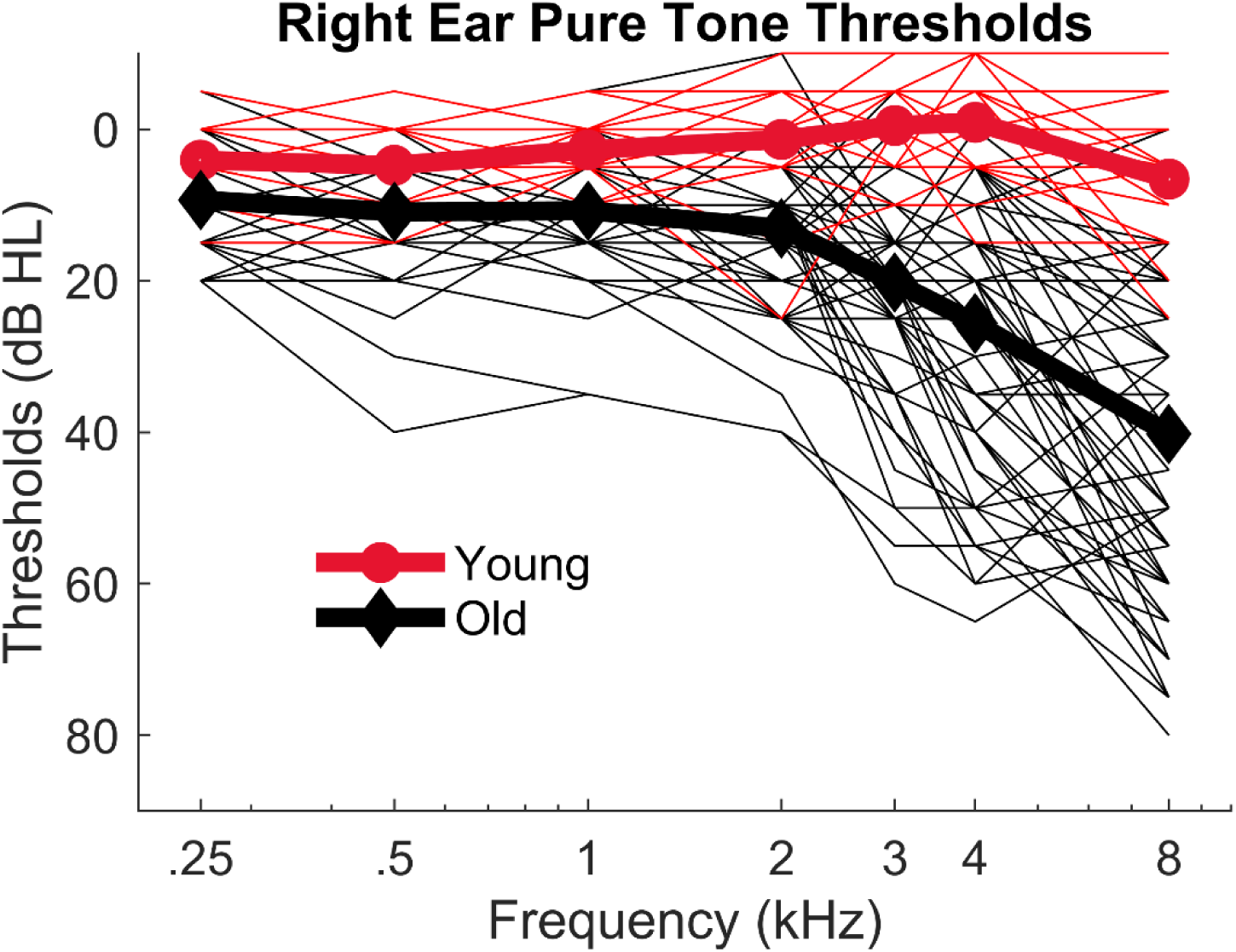
Audiometric thresholds of younger and older participants. Pure tone audiometric thresholds from younger (red circles) and older (black diamonds) participants. Individual audiometric thresholds are shown (thinner lines) with the average for each age group (thicker lines). Older adults have significantly higher thresholds than younger adults, particularly at higher test frequencies.

### Electrophysiological Measures of AN, Brainstem, and Cortical Function

To assess central gain, we examined the amplitude of the AN compound action potential (CAP) N1 response, auditory brainstem response (ABR) wave V, and cortical P1-N1 response evoked to a 100 dB pSPL click stimulus. To evaluate cortical activity in the AC and visual cortex (VC, serving as a control region), we used a distributed source model to estimate P1-N1 amplitudes within these regions of interest (ROI) as described below.

#### AN and ABR Acquisition and Analysis

CAPs were collected as part of several ongoing studies. Subsets of these CAP data have been reported elsewhere (Dias et al., 2024; Harris et al., 2021; McClaskey et al., 2018; Rumschlag et al., 2022), although their associations with ABR wave V and the cortical functional and structural characteristics described below are reported here for the first time. The CAP N1 and ABR wave V were elicited using 100 µs rectangular pulses (clicks) presented at a rate of 11.1/s with alternating polarity at 100 dB pSPL. 1100 trials, 550 of each polarity, were recorded. Stimuli were presented through an ER3C electrically-shielded insert earphone (Etymotic Technologies, IL, USA). CAP N1 responses were recorded using a tympanic membrane electrode (Sanibel, Denmark) in the test (right) ear, an inverting electrode placed on the contralateral (left) mastoid, and a low forehead grounding electrode. ABR wave V was simultaneously recorded using a high forehead active electrode, an inverting mastoid electrode (on the right mastoid), and a low forehead grounding electrode. All recordings were collected at a sampling rate of 20 kHz using a custom TDT (Tucker Davis Technologies, FL, USA) headstage connected to the bipolar channels of a Neuroscan SynAmpsRT amplifier in AC mode with 2010x gain (Compumedics, Canberra, Australia). Testing was done in an acoustically and electrically shielded room. Participants reclined in a chair and were encouraged to rest quietly for the duration of testing. Participants were allowed to sleep during CAP/ABR recording sessions. Continuous neural activity was analyzed off-line in MATLAB (MathWorks, Natick, MA) using the toolboxes EEGlab (Delorme and Makeig, 2004) and ERPLab (Lopez-Calderon and Luck, 2014). Continuous EEG signals were bandpass filtered between 0.150 kHz and 3.0 kHz. Stimulus triggers were delivered by TDT RPvdsEx and were shifted to account for the 1 ms delay introduced by the earphones and the 0.6 ms delay of the TDT digital-to-analog convertor. The filtered data were epoched from −2 to 11 ms and baseline corrected to a −2 ms to 0 ms prestimulus interval (McClaskey et al., 2018). Trials with peak deflections in excess of 45 µV were identified and rejected using an automatic epoch rejection algorithm that was visually inspected for accuracy. Epoched responses for the remaining trials were averaged.

CAP N1 and ABR wave V peak selection was performed by two independent and experienced reviewers and assessed for repeatability across multiple runs. Although only the 100 dB pSPL condition is reported here to match cortical levels, responses were collected from 70 to 110 dB pSPL to aid peak detection (e.g., Dias et al., 2024; Harris et al., 2022).

#### Cortical Response Acquisition

Cortical EEGs were recorded from a 64-channel Neuroscan QuickCap (international 10-20 system) connected to a SynAmps RT 64-channel Amplifier. Vertical electro-oculogram activity was recorded using bipolar electrodes placed above and below the left eye. A Krios digital scanner (NDI, Ontario, Canada) was acquired midway through data collection. Therefore, QuickCap electrodes were digitally mapped to the participant’s scalp for a sub-set of participants (n= 31). Of the 27 younger adults, 10 had mapped electrodes (7 female). Of the 48 older adults, 21 (12 female) had mapped electrodes. In participants missing the digitized map, a 64-channel Neuroscan QuickCap default electrode map, from the ICBM152 data set, was used. During EEG recordings, participants were allowed to recline in the same chair and room used for CAP and ABR recording. Participants were not allowed to sleep during cortical recordings and were instead instructed to read. Neural activity was recorded to a 100 µs 100 dB pSPL alternating polarity click with a 2000 ms inter-stimulus interval (20 ms jitter). This is the same stimulus used for CAP and ABR acquisition, but presented at a slower rate. Participants completed two sessions of 200 trials and the continuous EEG data were then processed offline using EEGLab and ERPLab. The recorded EEG data were down sampled to 0.5 kHz, bandpass filtered from 1 to 30 Hz, re-referenced to the average of all electrodes, and corrected for ocular artifacts using independent components analysis. Individual trials were then segmented into epochs around the click onset (−100 ms to +500 ms) and baseline corrected (−100 ms to 0 ms). Any epochs contaminated by peak-to-peak deflections in excess of 100 µV were rejected using an automatic artifact rejection algorithm.

#### Source-Constrained EEG Analyses

Source currents of the event related potentials (ERPs) were estimated using a cortical source estimation procedure in Brainstorm (Tadel et al., 2011). FreeSurfer (Fischl, 2012) was used to segment the T1 weighted image from each participant and create their individual boundary element model (BEM). Next, the MRI from each participant was co-registered with their respective EEG electrode map. While only a subset (n= 31) of participants had individual electrode maps, all participants had an MRI (see below) to create personalized BEM models.

MRIs from participants without mapped electrode locations were co-registered using the Brainstorm 64-channel Neuroscan QuickCap default electrode maps from the ICBM152 data set. Co-registration was performed using the nasion, left auricular point, right auricular point, anterior fissure, posterior fissure, and inter-hemispheric point as fiducials. The forward model was computed using the Symmetric BEM with OpenMEEG software with adaptive integration (Gramfort et al., 2010). Conductivities of the scalp, skull, and brain were set as 1, 0.0125, and 1, respectively. The noise covariance matrix for each participant was calculated from the pre-stimulus baseline. Source estimation was performed using the dynamic Statistical Parametric Mapping approach (Dale et al., 2000) with loose constraints (0.2), where minimum norm imaging was performed and then scaled with respect to the noise covariance, resulting in a map of z-scores. To localize responses to the left AC and left VC, scouts of both regions were defined from the S_temporal_transverse_L and G_cuneus_L areas of the Destrieux atlas (Destrieux et al., 2010), which closely overlap AC and VC, respectively. The final source-constrained EEG waveform for each region consisted of the averaged value from all vertex points within the scout.

Source-constrained ERP waveforms were analyzed using a custom MATLAB script. The approximate latencies of the first prominent negative peak (C1/N50), first prominent positive peak (P1), second prominent negative peak (N1), and second prominent positive peak (P2) were visually identified and recorded. These latencies were used to set the detection windows for the exact locations of the P1 and N1 peaks. P1 was defined as the maximum positive peak between C1/N50 and N1, and N1 was defined as the maximum negative peak between P1 and P2. The peak-to-peak P1-N1 amplitude deflection was calculated by computing the difference between the P1 amplitude and the N1 amplitude.

To determine if there was an effect of digitized or default EEG electrode map on observed P1-N1 deflections, we used linear mixed effects regression modeling (LMER). We found no significant difference in peak-to-peak amplitude between the digitized and default electrode maps for either AC P1-N1 [*β*= 0.109, t= -0.332, p= 0.741] or VC P1-N1 [*β*= 0.195, t= 0.802, p= 0.425]. Therefore, we did not distinguish between participants with and without mapped EEG electrode locations in our statistical analyses (see below).

### MRI and DKI Acquisition

Image acquisition was performed using a Seimens 3T Prisma^fit^ scanner with a 32-channel head coil (Siemens Medical, Erlangen, Germany). A T1 weighted image was obtained from all participants using a magnetization-prepared rapid gradient echo (MP2RAGE) with the following parameters: repetition time (TR)= 5000 ms, echo time (TE)= 2.98 ms, flip angle= 4°, 176 slices with a 256 × 256 matrix, slice thickness= 1.0 mm, and no slice gap, field of view (FOV)= 256 x 256 mm^2^. DKI scans were performed during the same session with b-values of 0, 1000, and 2000 s/mm^2^ applied in 64 directions, with the following parameters: TR= 3100, TE= 80.0 ms, FOV= 220 x 220 mm^2^, bandwidth= 1,136 Hz/pixel, phase partial Fourier encoding at 6/8, voxel size= 2.5 x 2.5 x 2.5 mm, slices= 50, slice thickness= 2.5 mm.

DKI data was processed using PyDesigner to compute diffusion parameters (Dhiman et al., 2021). Standard preprocessing was performed, consisting of Marchenko-Pastur principal component denoising, Gibbs ringing correction, eddy current correction, Echo Planar Imaging distortion correction, removal of non-brain tissue, gaussian smoothing, and Rician noise bias correction. Diffusion maps of computed fractional anisotropy (FA) and mean diffusivity (MD) were used for this study. FA values are generally thought to reflect the integrity of structures affecting water diffusion such as axonal membranes that affect the anisotropy, or directionality, of diffusion in cortical tissue (Van Hecke et al., 2016). Although FA can be computed in non-myelinated tissues, FA values are widely interpreted as reflecting general white matter integrity (Assaf and Pasternak, 2008). As white matter myelination decreases, FA values decline (Horsfield et al., 1998; Hüppi et al., 1998). MD values reflect overall diffusivity, regardless of direction, and are therefore highest in regions with no barriers to diffusion. Higher FA and lower MD are both associated with more microstructural integrity. Average FA and MD values were extracted from an ROI corresponding to the primary AC (TE 1.0 from the Julich Brain Atlas, Amunts et al., 2020) using the DSI studio software package (http://dsi-studio.labsolver.org). This area has a high degree of overlap with the area used for computed source-constrained ERPs for AC (see above). Whole brain FA and MD measures were calculated by averaging across the whole-brain FA or MD maps generated by PyDesigner using the mean function in the FSLmaths library of analysis tools (Jenkinson et al., 2012). Due to the limited field of view of the DKI scans, brainstem diffusion measures were not available for many of our participants, limiting our structural analyses to the cortex.

### SIN

We measured SIN using the Quick Speech-in-Noise Test (QuickSIN; Etymotic Technologies) (Killion et al., 2004). QuickSIN data were collected from all participants. The QuickSIN test consists of five lists of six low-probability sentences, each with five keywords per sentence, totaling 30 keywords in each list. Sentences were presented in a four-talker babble. Sentences were presented binaurally using TDH-39 headphones using a combination of an Onkyo Compact Disk Player (Onkyo, Osaka, Japan) and an Interacoustics AA222 Audio Traveler (Interacoustics, Middelfart, Denmark). For each list, the six sentences are presented in a progressively decreasing signal-to-noise ratio (SNR) from 25 to 0 dB in 5 dB steps, with sentences presented at 70 dB HL and the background noise level varying. The average number of keywords per sentence correctly identified at each SNR (25, 20, 25, 20, 5, 0 dB) was calculated. Then, we summed the number of correctly identified keywords to arrive at a total correct score out of a possible 30. QuickSIN performance is reported as SNR loss, calculated as 25.5 minus the total keywords correct (Killion et al., 2004, Etymotic Research). Less SNR loss indicates better SIN performance.

### Experimental Design and Statistical Analysis

Statistical analyses were performed in R version 4.3.1 (R Core Team, 2021). General linear modeling and generalized linear mixed-effects modeling were performed using the lme4 package (Bates et al., 2015). Sex differences in our variables were not hypothesized, and preliminary analyses confirmed there were no significant sex differences in response amplitudes, diffusion metrics, or SIN recognition, nor did including sex as a covariate in our statistical models improve model fit (p>0.05).

#### Progression of Central Gain Through the Levels of the Auditory Pathway

Animal studies have shown that deficits in inhibitory control at the brainstem contribute to central gain (Palombi et al., 2001; Keine et al., 2016; Ono and Ito, 2024) and our prior work has confirmed a similar effect in humans (Rumschlag et al., 2022). Additionally, our work (Harris et al., 2022) and others (Overton and Recanzone, 2016; Dobri and Ross, 2021; Xue et al., 2023) suggest that age-related deficits in structure and function at the AC may contribute to central gain in older adults. We hypothesized that age-related loss of afferent auditory activity contributes to central gain, with gain occurring at the brainstem and further amplified at the cortex. To test this, we examined gain using a multivariate approach that tested for age group differences in AN (CAP N1), brainstem (ABR wave V), and source-constrained AC activity (P1-N1). We used source-constrained VC activity (P1-N1) as a cortical control. Illustrated in Figure 2, we hypothesized that age-related deficits in AN activity would predict a recovery of response amplitudes – the absence of an age-group difference – at the auditory brainstem and an amplified response at the AC in older adults, relative to responses from younger adults.

**Figure 2:**
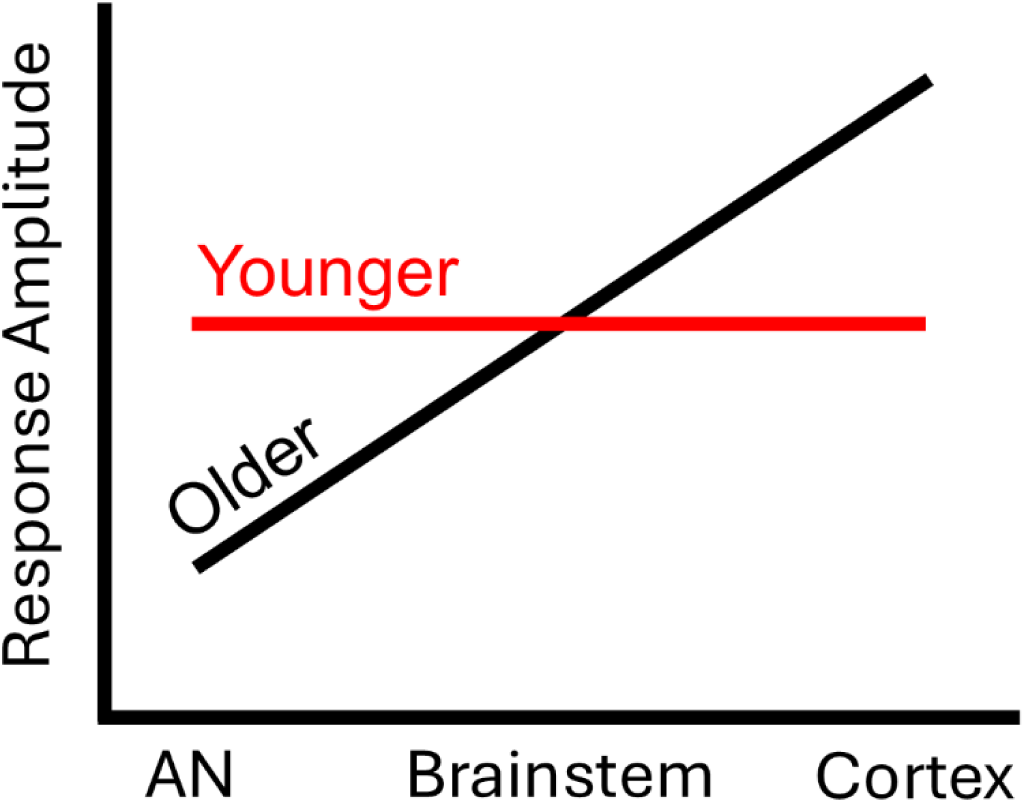
Hypothesized progression of central gain through the levels of the auditory pathway. We hypothesize that there is progression of central gain in older adults from the auditory brainstem to the auditory cortex. Illustrated here, we predict that auditory nerve (AN) deficits in older adults (relative to the responses of younger adults) predict a recovery of response amplitudes at the auditory brainstem (a lack of age-group differences) and amplified responses at the auditory cortex.

To test this progression of central gain through the auditory system, we used a general linear model approach. Consistent with our hypothesis, we predicted that older adults, compared to younger adults, would have AN responses (CAP N1 amplitudes) that were smaller, similar responses at the auditory brainstem (ABR wave V amplitudes), and larger responses in AC (AC P1-N1 amplitudes). To determine the extent to which the relationships we observed with the AC are specific to the AC, we also examined relationships with responses measured in VC (see “Source-constrained EEG analysis pipeline”). The units of measure for source-constrained ERPs are arbitrary; this is a byproduct of source reconstruction. Response amplitudes can also vary systematically from one level of the auditory system to another. As such, for these analyses testing the progression of gain, amplitudes were z-scored at each level of the auditory pathway (CAP N1, ABR wave V, and source-constrained ERP P1-N1) and visual cortex (source-constrained ERP P1-N1) across age groups. CAP N1 amplitudes were z-scored after being converted to absolute values so that larger (more negative) responses would correspond to positive z-scores. By z-scoring response amplitudes in this way, we can compare age-group differences in response amplitude at different levels of the auditory pathway. In our statistical models, z-scored amplitudes were the outcome variable, with level of the auditory system (CAP N1, ABR wave V, AC P1-N1, or VC P1-N1) and age group (younger or older) as categorical predictors. Participant was included as a random effects factor. This was followed by hypothesis-driven planned linear regression models to examine age-group differences in response amplitudes at each level of the auditory pathway. We also tested if average audiometric thresholds (PTA) were predictive of response amplitudes.

#### AN Activity and Cortical Microstructure

Although mixed, there is some evidence to suggest that hearing loss may be associated with changes in cortical diffusion metrics, including FA and MD (Tarabichi et al., 2018; Husain et al., 2011; but see Profant et al., 2014). Age effects on cortical diffusion metrics are well-established, yet highly variable across the cortex (Zhao et al., 2015; Gray et al., 2020; Schilling et al., 2023).

Additionally, AN responses are highly variable in both younger and older adults, and aging is known to contribute to smaller AN response amplitudes (Konrad-Martin et al., 2012; Grose et al., 2019; Harris et al., 2022; Rumschlag et al., 2022). Because of these relationships between age, AN function, and diffusion metrics of cortical microstructure, we examined the extent to which afferent input (CAP N1 amplitudes) was predictive of diffusion metrics (FA and MD) in the AC of younger and older adults. We hypothesized that if afferent loss exacerbates age-related cortical degeneration, then CAP N1 deficits would predict lower FA and higher MD in the AC of older adults. We tested this hypothesis using linear regression models to examine the extent to which afferent auditory input (CAP N1 amplitudes) predicted diffusion metrics calculated in AC, accounting for age-group differences and whole-brain diffusion metrics.

#### AC Microstructure, Cortical Responses, and Central Gain

To identify cortical structure-function associations, and to test the hypothesis that poorer microstructural integrity may result in increased central gain, similar to associations found in animal models (Borges et al., 2023), we used linear regression models to test the degree to which AC metrics for microstructural integrity (FA and MD) predicted cortical response amplitudes (AC P1-N1) in younger and older adults. Due to the high degree of collinearity between these diffusion metrics, separate models were run for FA and MD.

If FA or MD was found to be a significant predictor of AC P1-N1, then the whole brain average of the specific metric (FA or MD) was incorporated into the model as a control to test the degree to which the observed relationship is specific to AC.

Building upon these analyses, we used linear regression models to test the degree to which AC FA and MD were significant predictors of auditory cortical gain AC gain in older adults. AC gain in older adults was quantified using the same method described in Harris et al. (2022). In short, CAP N1 amplitudes predict cortical P1-N1 amplitudes in younger adults. The parameters of the linear model derived from this relationship in younger adults were used to predict the expected cortical response in each older adult, based on their CAP N1 amplitude, using the predict.lm function from the R package “stats” (R Core Team, 2021). To determine if AC gain was evident in our older adults, we used a paired-samples t-test to test if observed auditory cortical responses were larger than those predicted from the CAP N1 amplitude. To quantify the AC gain in each participant, the predicted auditory cortical response was subtracted from the observed auditory cortical response (Harris et al., 2022).

### AC Microstructure and SIN in Younger and Older Adults

Several factors are thought to contribute to SIN difficulties in older adults. Our prior study found that greater central gain was associated with poorer SIN recognition in older adults (Harris et al., 2022). Myelin integrity is hypothesized to contribute to temporal precision in the auditory system (Zeng et al., 2005; Long et al., 2018; Harris et al., 2021), which is crucial for SIN recognition. Therefore, we examined the extent to which deficits in AC diffusion metrics (FA and MD), which are thought to relate to myelin integrity, predict SIN recognition in older adults. We used a model testing approach to test the hypothesis that structural deficits in the AC predict poorer SIN recognition. AC diffusion metrics, whole brain diffusion metrics, and PTA were included as predictors, with SIN as the outcome variable. Due to collinearity between FA and MD, separate models were run for FA and MD. Separate models were tested in younger and older adults.

## Results

### Central Gain Propagates Through the Auditory Pathway

Average waveforms for click-evoked CAPs, ABRs, AC P1-N1, and VC P1-N1 are shown in Figure 3. An LMER model examining the extent to which age-group and level (AN, brainstem, cortex) of the auditory pathway predict response amplitudes found a significant interaction between age group and level (Table 1). PTA was not a significant predictor of response amplitude across levels of the auditory system and VC [*β*= -0.107, t= 0.098, p= 0.616] and incorporating PTA into our model did not improve model fit [χ²(1)= 1.675, p= 0.196], which is consistent with prior studies showing that hearing thresholds are not predictive of suprathreshold averaged responses (Parthasarathy and Kujawa, 2018; Harris et al., 2022; Rumschlag et al., 2022). Subsequent LM models examining age-group differences in response amplitude at each level of the auditory pathway are reported in Table 1. Consistent with our prior reports (Harris et al., 2021, 2022), age group was a significant predictor of AN response amplitudes, with older adults exhibiting significantly smaller AN responses than younger adults [Younger: mean= 0.350, SD= 0.796; Older: mean= -0.168, SD= 1.061]. ABR wave V amplitudes did not significantly differ between younger and older adults [Younger: mean= -0.224, SD= 0.960; Older: mean= 0.122, SD= 1.022]. AC P1-N1 amplitudes were significantly larger in older adults than in younger adults [Younger: mean= -0.401, SD= 0.828]; Older: mean= 0.190, SD= -0.401]. VC response amplitudes were not significantly different between younger and older adults [Younger: mean= -0.026, SD= 1.075; Older: mean= 0.015, SD= 0.977]. The preservation of VC amplitudes may reflect cross-sensory influences in older adults (Glick and Sharma, 2017) or may be due to spatial leakage across source-constrained ROIs. Individual response amplitudes are reported in Figure 4. The results support our hypothesis that central gain propagates through the levels of the auditory system, showing that older adults with age-related deficits in AN activity can exhibit gain at the auditory brainstem (a recovery of responses amplitudes relative to age-related deficits at the AN) and a progressive increase in gain at the AC (amplified AC responses relative to younger adults).

**Figure 3:**
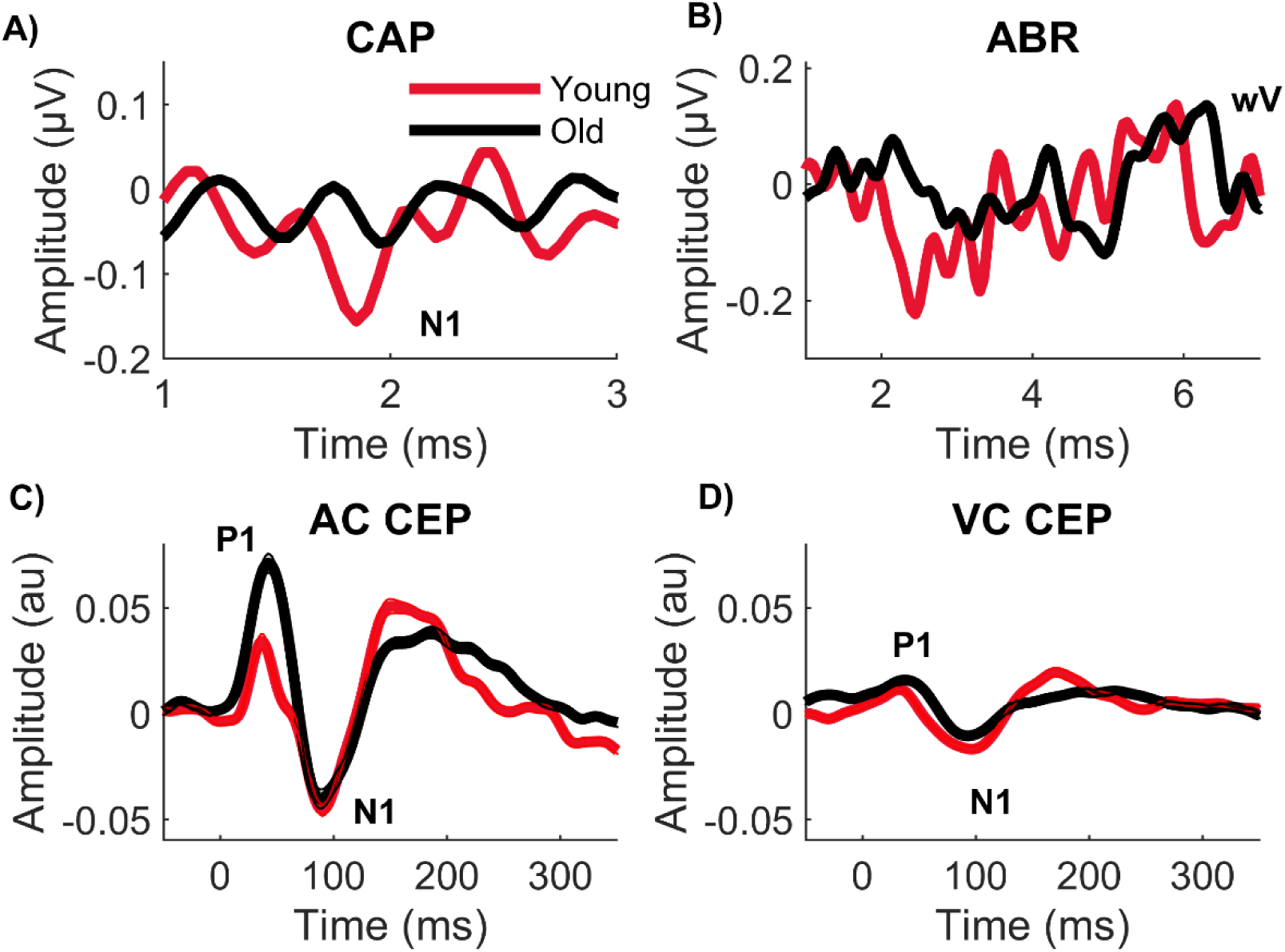
Average waveforms for younger and older adults. A-D) Average waveforms in response to 100 dB pSPL clicks presented to the right ear in older and younger adults. A) Compound action potential (CAP) waveforms. N1, corresponding to the auditory nerve response, is marked. B) Auditory brainstem response (ABR) waveforms. Wave V (wV), corresponding to the brainstem response, is marked. C) Cortical event-related potential (ERP) waveforms source-constrained to the auditory cortex (AC) with standard error of the mean. D) ERP waveforms source-constrained to the visual cortex (VC) with standard error of the mean. P100 (P1) and N100 (N1) are marked. “au”= arbitrary units.

**Figure 4:**
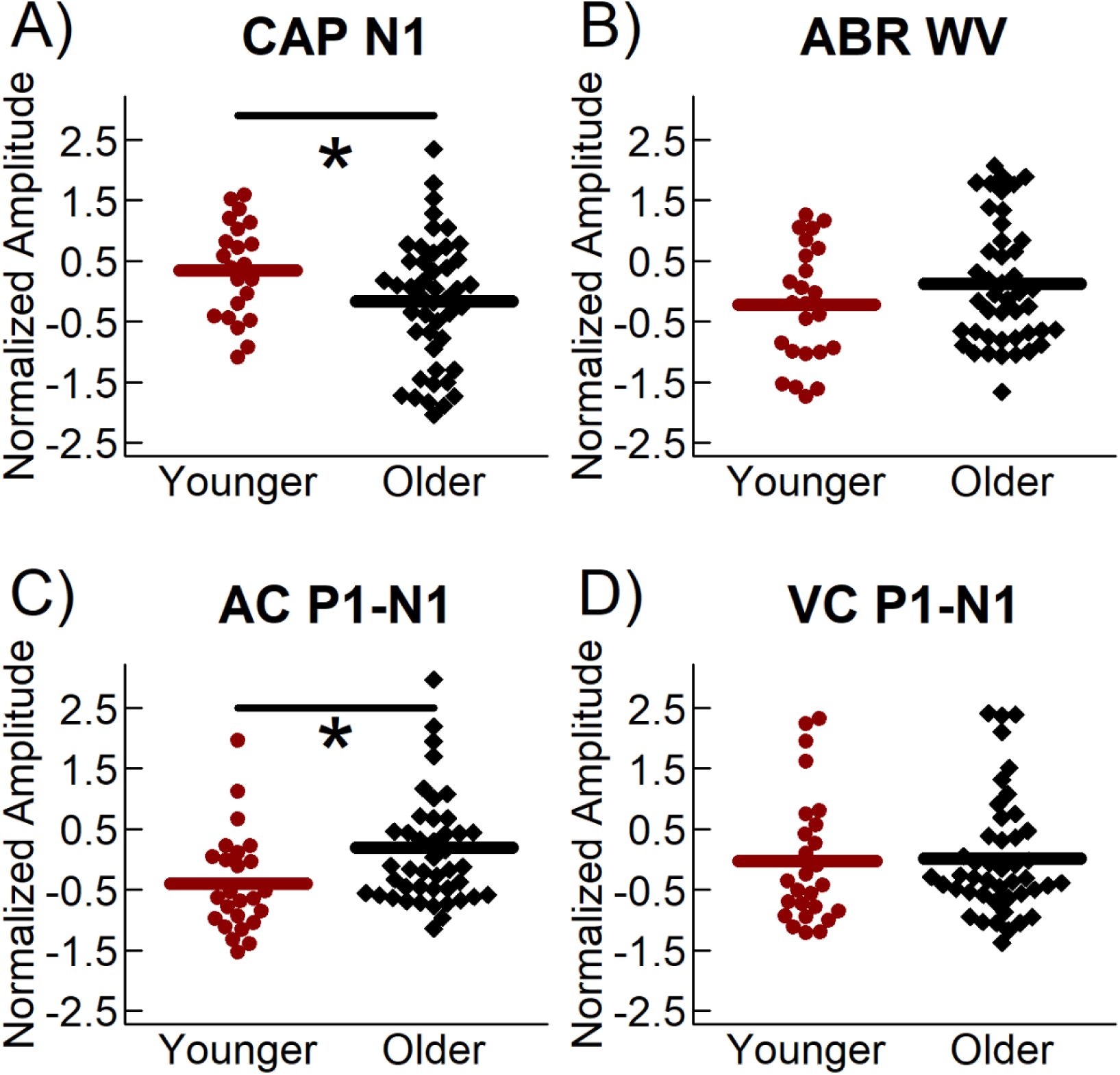
Older adults exhibit progression of central gain. Data points represent the individual peak amplitudes for younger (red circles) and older (black diamonds) participants plotted as a function of age group and location. Means marked. Amplitudes are represented as a z-score. A) Compound action potential (CAP) N1 amplitude, represented as the absolute value. B) ABR wave V amplitude. C) Auditory cortex (AC) P1-N1 amplitude. D) Visual cortex (VC) P1-N1. Compared to younger adults, older adults exhibit a significantly smaller N1 (p= 0.004) and a significantly larger AC P1-N1 (p= 0.039) with no difference in wave V amplitude or VC P1-N1. * <0.05.

**Table 1:** Auditory-evoked response amplitudes: Age-group differences across levels of the auditory system.

| Amplitudes | df | F | p |  |
| --- | --- | --- | --- | --- |
| Level | (3, 266) | 0.003 | 0.999 |  |
| Age Group | (1, 266) | 0.269 | 0.604 |  |
| Level x Age | (3, 226) | 3.317 | 0.020 | * |

| Planned Linear Models | B | SEB | $\beta$ | t | p | |
| --- | --- | --- | --- | --- | --- | --- |
| CAP N1 amplitude |  |  |  |  |  |  |
| Age Group | -0.518 | 0.255 | -0.242 | -2.028 | 0.047 | * |
| ABR wave V amplitude |  |  |  |  |  |  |
| Age Group | 0.347 | 0.254 | 0.166 | 1.364 | 0.177 |  |
| AC P1-N1 amplitude |  |  |  |  |  |  |
| Age Group | 0.040 | 0.019 | 0.248 | 2.107 | 0.039 | * |
| VC P1-N1 amplitude |  |  |  |  |  |  |
| Age Group | -0.079 | 0.257 | -0.038 | -0.307 | 0.760 |  |

\* $p < 0.05$ , \*\* $p < 0.01$ , \*\*\* $p < 0.001$ . CAP= compound action potential. ABR= auditory brainstem response. AC= auditory cortex. VC= visual cortex.

### AN Activity Predicts AC Microstructure

AC FA [*β*= -0.312, t= -2.767, p= 0.007] and whole brain FA [*β*= -0.405, t= -3.730, p<0.001] were significantly different between age groups, with older adults exhibiting lower FA than younger adults [AC: Younger, mean= 0.195, SD= 0.047; Older, mean= 0.167, SD= 0.037. Whole brain: Younger, mean= 0.311, SD= 0.021; Older, mean= 0.289, SD= 0.024]. Similarly, both AC MD [*β*= 0.498, t= 4.806, p<0.001] and whole brain MD [*β*= 0.603, t= 6.330, p<0.001] were significantly increased in older adults [AC: Younger, mean= 1.112, SD= 0.149; Older, mean= 1.571, SD= 0.473. Whole brain: Younger, mean= 1.319, SD= 0.139; Older, mean= 1.533, SD= 0.137]. Taken together these results show microstructural deficits in the AC and whole brain of older adults.

CAP N1 amplitudes were a significant predictor of AC FA, with larger CAP N1 amplitudes associated with higher AC FA (Table 2, Figure 5), even when controlling for both age and whole brain FA, supporting our hypothesis that reduced afferent auditory input affects AC microstructure. AC FA was not correlated with whole brain FA [*β*= 0.091, t= 0.774, p= 0.442]. The results suggest that reduced AN responses affect AC FA beyond general age-related decreases in FA observed across the cortex. CAP N1 amplitudes did not predict AC MD [*β*= -0.114, t= 0.708, p= 0.213]. AC MD and whole brain MD were significantly positively related [*β*= 0.715, t= 8.552, p<0.001], suggesting that age-related deficits in MD are similar across the cortex and are not influenced by AN function.

**Figure 5:**
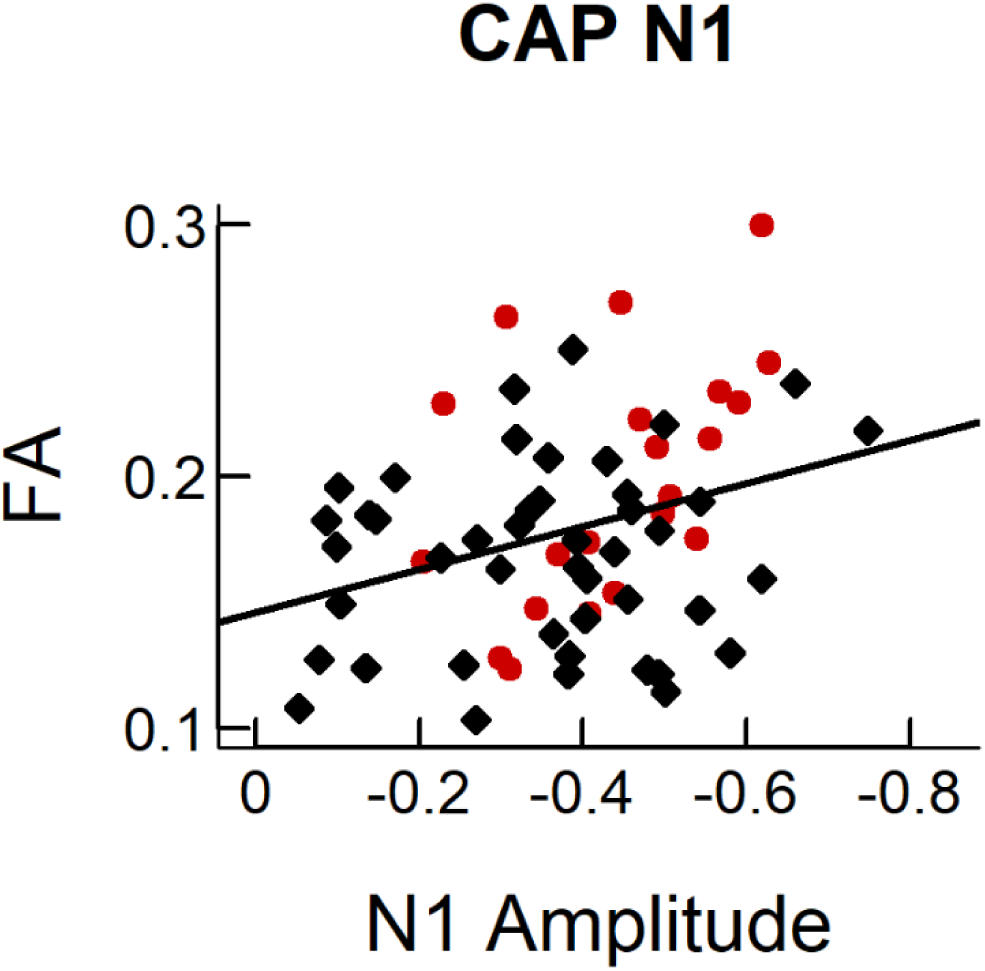
Larger AN responses predict higher AC FA in older adults. Data are the individual data points from older (black diamond) and young (red circles) participants. Compound action potential (CAP) N1 amplitudes, reflecting the auditory nerve (AN) response, are a significant predictor of fractional anisotropy (FA) in the auditory cortex (AC) across age groups. Solid black lines indicated a significant relationship between variables.

**Table 2:** AC FA- effects of age and CAP N1 amplitude.

| **Factors** | **B** | **SE<sub>B</sub>** | **$\beta$** | **t** | **p** |  |
| --- | --- | --- | --- | --- | --- | --- |
| Age Group | -0.024 | 0.011 | -0.269 | -2.063 | 0.043 | * |
| CAP N1 amplitude | -0.068 | 0.032 | -0.253 | -2.114 | 0.039 | * |
| Whole-brain FA | -0.049 | 0.214 | -0.014 | -0.116 | 0.908 |  |

\*p<0.05, \*\*p<0.01. CAP= compound action potential. FA= fractional anisotropy.

The results support our hypothesis that age-related AN deficits can predict microstructural deficits in the auditory cortices of older adults (FA). Importantly, this relationship between afferent auditory input and AC microstructure holds true even after accounting for age and age-related deficits in whole brain microstructure, emphasizing the influence AN function may have on the structural integrity of the AC.

### AC Microstructure Predicts Cortical Responses and Central Gain in Older Adults

Table 3 reports the results of the linear regression model testing our hypothesis that AC microstructure predicts AC P1-N1 amplitudes. AC FA was a significant predictor of AC P1-N1 amplitude across age-group, while age-group and whole brain FA were not significant predictors. However, there was a significant interaction between age group and AC FA. Thus, we used separate linear models to examine associations between AC FA and response amplitudes in younger and older adults, controlling for whole brain FA. In younger adults, neither AC FA nor whole brain FA predicted cortical AC P1-N1 response amplitudes (Table 3, Model 1). In contrast, lower AC FA in older adults predicted larger AC P1-N1 response amplitudes, even after controlling for whole brain FA (Table 3, Model 2). When we examined the same associations with AC MD, we found that AC MD was not a significant predictor of AC P1-N1 amplitudes [*β*=0.022, t=0.9278, p=0.357].

**Table 3:**
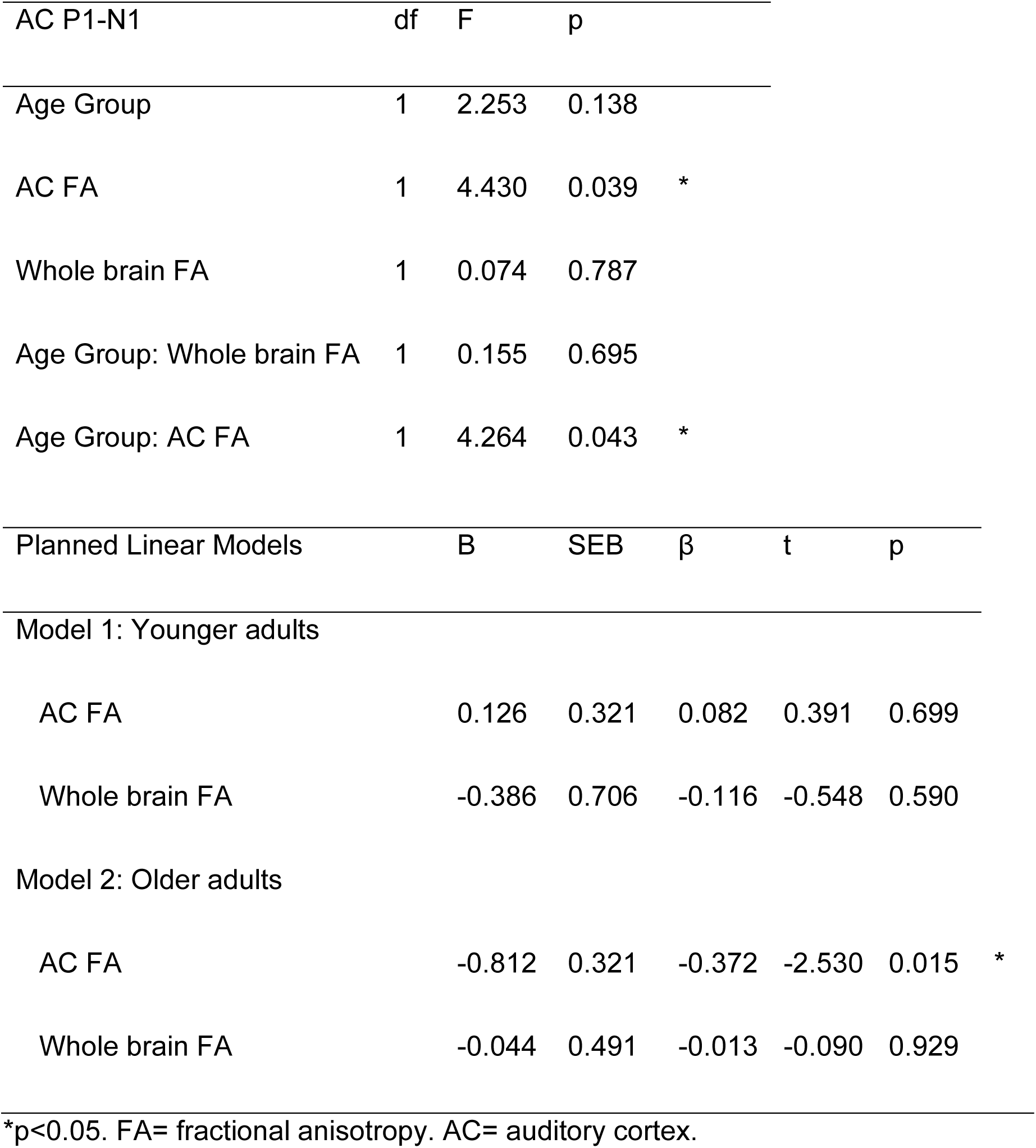
AC P1-N1- effects of age and FA.

| AC P1-N1 | df | F | p |  |
| --- | --- | --- | --- | --- |
| Age Group | 1 | 2.253 | 0.138 |  |
| AC FA | 1 | 4.430 | 0.039 | * |
| Whole brain FA | 1 | 0.074 | 0.787 |  |
| Age Group: Whole brain FA | 1 | 0.155 | 0.695 |  |
| Age Group: AC FA | 1 | 4.264 | 0.043 | * |

| Planned Linear Models | B | SEB | $\beta$ | t | p | | |
| --- | --- | --- | --- | --- | --- | --- | --- |
| Model 1: Younger adults |  |  |  |  |  |  |  |
| AC FA | 0.126 | 0.321 | 0.082 | 0.391 | 0.699 |  |  |
| Whole brain FA | -0.386 | 0.706 | -0.116 | -0.548 | 0.590 |  |  |
| Model 2: Older adults |  |  |  |  |  |  |  |
| AC FA | -0.812 | 0.321 | -0.372 | -2.530 | 0.015 | * |  |
| Whole brain FA | -0.044 | 0.491 | -0.013 | -0.090 | 0.929 |  |  |

\*p<0.05. FA= fractional anisotropy. AC= auditory cortex.

The relationship between AN response amplitudes (CAP N1) and AC FA and the relationship between AC response amplitude (AC P1-N1) and AC FA occur in opposite directions in older adults. Smaller AN responses predicted lower AC FA, but lower AC FA predicted larger AC P1-N1 amplitudes (Figure 6). These results are broadly consistent with models of central gain and age-related central nervous system degeneration. Although CAP N1 amplitudes predict AC FA across age groups, AC FA only predicts cortical response amplitudes in older adults. This may be due to age-related degeneration in the AC coupled with central compensation for reduced afferent auditory input.

**Figure 6:**
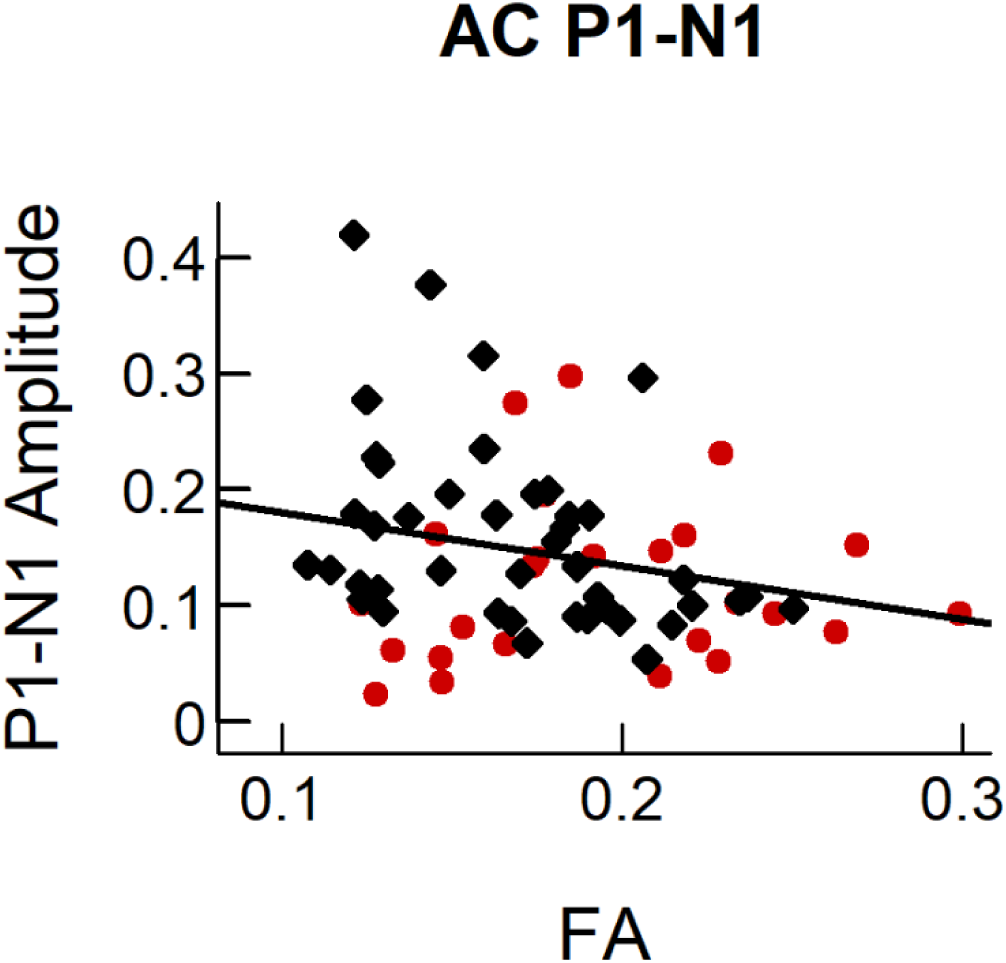
Lower AC FA predicts larger AC response amplitudes in older adults. Data are the individual data points from older (black diamond) and young (red circles) participants. Responses from the auditory cortex (AC) P1-N1 amplitudes decrease with increasing fractional anisotropy (FA) in the AC across age groups. The solid black line indicates a significant relationship between variables.

We quantified central gain as the difference between the AC P1-N1 response amplitudes observed in our older adults and the AC P1-N1 response amplitudes predicted from their CAP N1 response amplitudes (using parameters for the relationship between CAP N1 and AC P1-N1 measured in younger adults). This metric of central gain represents the degree to which AC response amplitudes are larger than would be expected from afferent input (Harris et al., 2022). We used this metric of central gain to test our hypothesis that lower AC FA, which predicted larger AC P1-N1 response amplitudes (see above), also predicts more AC gain in older adults. We note that, similar to our prior study, older adults exhibited significant central gain, such that observed AC P1-N1 responses were significantly larger than those predicted from their CAP N1 response amplitudes [p<0.001]. Our linear model found that lower AC FA predicted more AC gain in older adults [*β*= -0.402, t= -2.596, p= 0.014]. AC MD did not predict AC gain [*β*= 0.079, t= 0.460, p= 0.627]. The results support our hypothesis that deficits in AC microstructure (lower FA) predict central gain in older adults.

### AC FA and MD are Predictive of SIN Recognition in Older Adults

Consistent with our hypothesis that myelin deficits contribute to poorer SIN recognition in older adults, our linear regression model showed that lower AC FA predicted more SNR loss (poor SIN recognition) in older adults (Figure 7.A) [*β*= -0.372, t= -2.387, p= 0.022]. Adding whole brain FA to this model did not significantly improve model fit [χ²(1)= 0.033, p= 0.858], nor was whole brain FA a significant predictor of SNR loss [*β*= -0.124, t= -0.948, p= 0.347]. Similarly, adding PTA did not improve model fit [χ²(1)= 1.956, p= 0.151] and PTA was not a significant predictor of SNR loss [*β*= 0.178, t= 1.111, p= 0.263]. AC FA was not a significant predictor of SNR loss in younger adults [*β*= -0.196, t= -0.913, p= 0.371]. Lower AC MD predicted better SIN recognition in older adults (Figure 7B) [*β*= 0.401, t= 2.550, p= 0.015]. Adding PTA did not improve model fit [χ²(1)= 0.706, p= 0.407], and was not a significant predictor of SIN in this model [*β*= 0.059, t= 0.840, p= 0.731]. AC MD was not a significant predictor for SIN in younger adults [*β*= 0.332, t= 1.610, p= 0.122]. The results support our hypothesis that deficits in AC microstructure, consistent with a loss of myelin, predict poor SIN recognition in older adults.

**Figure 7:**
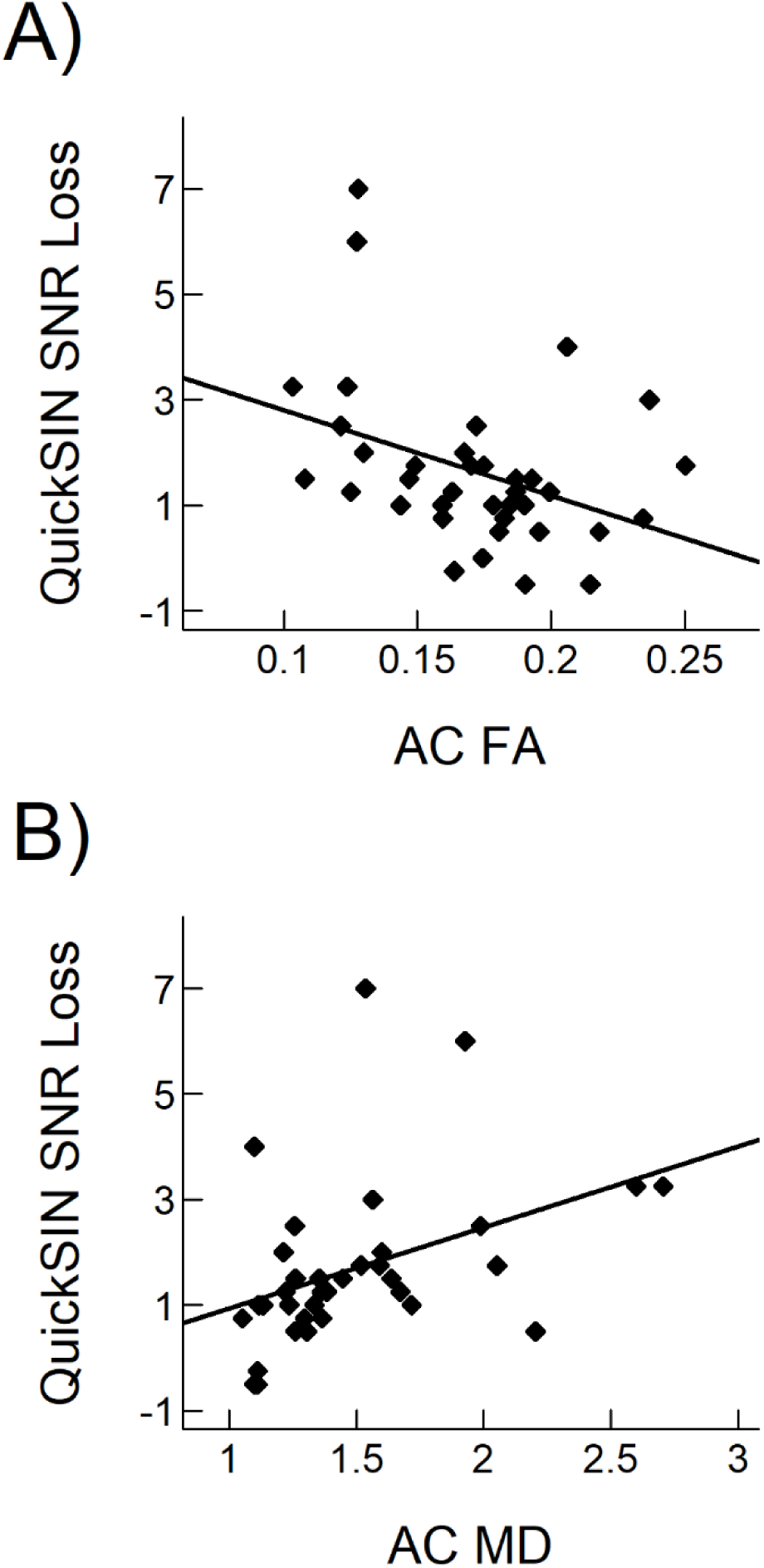
Poorer microstructural integrity in the AC predicts better SIN recognition in older adults. All data are from older adults. A) Higher auditory cortex (AC) fractional anisotropy (FA) is predictive of better speech-in-noise (SIN) recognition. B) Higher AC mean diffusivity (MD) is predictive of worse SIN recognition. The solid black line indicates a significant relationship between variables.

## Discussion

The underlying mechanisms and spatial progression of age-related auditory central gain and the contribution to SIN recognition are not well understood. Evidence from human and animal models shows that a loss of afferent auditory input results in compensatory central gain and structural changes in multiple regions of the auditory system, including the brainstem and cortex (Chambers et al., 2016; Harris et al., 2022; Rumschlag et al., 2022; Hutchison et al., 2023). In this study, we demonstrated that 1) older adults exhibit a spatial progression of central gain through the levels of the auditory system, with evidence for gain at the auditory brainstem and additional gain at the AC; 2) in older adults, reduced afferent auditory input predicted lower AC white matter integrity (lower AC FA); 3) AC FA deficits predicted larger auditory-evoked responses and more central gain in the AC of older adults; and 4) reduced AC microstructural integrity (lower AC FA and higher AC MD) is associated with poorer SIN recognition in older adults.

### Central Gain in the Brainstem and Cortex

Our results expand on previous studies showing age-related central gain in both the auditory brainstem (Grose et al., 2019; Parthasarathy et al., 2019; Johannesen and Lopez-Poveda, 2021; Rumschlag et al., 2022) and AC (Alain et al., 2022; Harris et al., 2022). Findings from mice have shown that cochlear denervation results in relatively more central gain in the AC compared to the inferior colliculus (Chambers et al., 2016). Translating these results to older adults, we demonstrated that the magnitude of central gain, when compared to younger adults, is larger in the AC than in the brainstem. This is the first study to show that central gain progressively increases through the levels of the human auditory system, with amplification of responses at the brainstem followed by further amplification at the AC. Although the mechanisms underlying central gain are not fully known, work from animal models and our current study have linked gain to myelin degeneration or hypomyelination in the AC (Borges et al., 2023). Similarly, our prior work and others have linked central gain to reduced inhibition in the brainstem and AC (Keine et al., 2016; Salvi et al., 2017; Harris et al., 2022). Further studies are needed to better understand how altered signaling and central gain in the lower levels of the auditory system (e.g., inferior colliculus) might interact with changes at the cortex to exacerbate gain at higher levels of the auditory system.

### AN Activity and AC Microstructural Complexity

Our results support the notion that reduced afferent input can result in a deficit in myelin integrity in the central auditory system. Myelin is integral for ensuring the synchronous neural responses needed for generation of the CAP. Degeneration of peripheral myelin is known to negatively affect AN responses in human and animal models (Panganiban 2022; Chiappa et al., 1980; Marangos, 1996). For the current investigation we used diffusion metrics, FA and MD, that are hypothesized to reflect the microstructural integrity of underlying tissue, including myelin. Smaller FA and larger MD values are generally interpreted as reflecting disruptions in tissue structure. While MD indicates the rate of water diffusion through the tissue and can serve as a general metric of tissue integrity, FA indicates the directionality of diffusion and may reflect white-matter fiber integrity more specifically, including myelin integrity. In humans, hearing loss has been shown to relate to both FA and MD in the inferior colliculus and AC (Husain et al., 2011; Profant et al., 2014; Ma et al., 2016), although these relationships are inconsistent across studies and interactions with age were often not considered (Tarabichi et al., 2018). We found that age-related deficits in afferent auditory activity may contribute to lower AC FA in older adults and potentially exacerbate age-related microstructural declines.

### Potential Role of Myelin in Central Gain

Our finding that AC FA – a proxy measure of white matter myelin integrity – is a significant predictor of central gain is consistent with evidence from animal models that hypomyelination in the auditory system may result in hyperexcitation (Borges et al., 2023). Our results also complement our previous work finding a link between AC gain and gamma-aminobutyric acid (GABA) (Harris et al., 2022). Together, these findings suggest that age-related changes in myelin and GABA, perhaps as a result of age-related AN deficits, can account for the central gain observed in older adults. GABA can directly affect cortical myelin, such that modulation of GABA can increase the population of mature oligodendrocytes and improve myelination (Zonouzi et al., 2015; Shaw et al., 2019), and FA measures (Cisneros-Mejorado et al., 2020, 2024) in animal models.

However, only a single study thus far has examined the relationship between GABA and diffusion metrics of white matter integrity (FA), but only within younger adults (Ziminski et al., 2023). More work is needed to understand the link between GABA and myelin, and how age-related changes in these biological factors can contribute to central gain and speech recognition deficits.

### Associations with SIN

As we stated earlier, myelin integrity is essential for the efficient and accurate encoding of complex stimuli, like speech (Zeng et al., 2005). Previous research has shown the importance of white matter tracts in speech recognition (Shekari and Nozari, 2023), including SIN recognition in older adults (Perron et al., 2021). Despite older adults exhibiting greater difficulties perceiving SIN, studies on the effect of age on FA and MD measures in the auditory system are mixed, with some showing a significant reduction in white matter integrity with age (Husain et al., 2011; Ma et al., 2016) while others showing no effect of age (Profant et al., 2014; Koops et al., 2021). Our current results support previous findings that preserved microstructural integrity contributes to better SIN recognition in older adults.

### Myelin as a Potential Therapeutic Target for SIN Difficulties

Understanding the relationships between afferent loss, central gain, myelin integrity, and speech recognition could lead to new targets for intervention. Our results show that AC microstructural integrity predicts SIN recognition in older adults. Myelin can change throughout the lifespan in response to experience (Osso and Hughes, 2024). Social isolation (Liu et al., 2016) or acute hearing loss (Kurioka et al., 2021) can result in myelin deficits, while learning can increase oligodendrogenesis and myelination (de Villers-Sidani et al., 2010; Hughes et al., 2018; Bacmeister et al., 2020). Evidence from human models also suggests a connection between learning and white matter modulation (Scholz et al., 2009). Although the exact mechanisms by which myelin integrity influences SIN recognition are relatively unknown, the ability of myelin to dynamically change throughout the lifespan, coupled with our findings that AN function may modulate AC microstructure, suggests a potential novel target to treat age-related deficits in SIN.

### Potential Limitations

The current study used diffusion imaging to assess white matter integrity. DKI metrics for white matter microstructure (FA and MD) are associated with myelin, but can be influenced by other factors (e.g., fiber dispersion, inflammation) (Beaulieu, 2002; Takahashi et al., 2002; Jones et al., 2013; Jiang et al., 2021). Novel techniques for imaging myelin, including myelin water imaging, may improve our understanding of the mechanisms contributing to cortical degeneration and central gain observed in the current study. Another limitation is that, due to field of view constraints, we were not able to examine the relationship between brainstem white matter microstructure and central gain. Future studies that examine myelin throughout the auditory system are needed to address these remaining knowledge gaps.

## Summary and conclusions

Our results demonstrate that older adults exhibit progressive central gain at the level of the auditory brainstem and cortex. In older adults, reduced afferent auditory input predicted reduced AC white matter integrity, which then predicted AC gain. Deficits in microstructural integrity predict poorer SIN recognition in older adults. Together, these results suggest that loss of afferent auditory input in older adults triggers maladaptive plasticity in the auditory system, resulting in progressive central gain and altered cortical microstructure that may contribute to poorer SIN recognition. Several gaps in knowledge remain and future studies using myelin-specific metrics are needed to further our understanding of the relationship between auditory afferent loss, central gain, and AC structural integrity.

## Acknowledgements and Funding Sources

This work was supported (in part) by grants from the National Institute on Deafness and Other Communication Disorders (NIDCD) (P50 DC000422, R01 DC017619, R01 DC021064, R21 DC020559, and T32 DC014435) and the Hearing Health Foundation. The project received support from the South Carolina Clinical and Translational Research (SCTR) Institute with an academic home at the Medical University of South Carolina, National Institute of Health/National Center for Research Resources (NIH/NCRR) Grant number UL1 TR001450. This investigation was conducted in a facility constructed with support from Research Facilities Improvement Program Grant Number C06 RR14516 from the NIH/NCRR.

The authors declare no competing financial interests or conflicts of interest.

